# Generalizability of Normative Models of Brain Morphometry Across Distinct Ethnoracial Groups

**DOI:** 10.1101/2024.10.14.618114

**Authors:** Ruiyang Ge, Yuetong Yu, Faye New, Shalaila S Haas, Nicole Sanford, Kevin Yu, Julian Camillo Becerra Leon, Guoyuan Yang, Jia-Hong Gao, Kiyotaka Nemoto, Masaki Fukunaga, Junya Matsumoto, Ryota Hashimoto, Neda Jahanshad, Paul M Thompson, Sophia Frangou

## Abstract

Normative modeling of brain morphometric data can inform about the clinical significance of deviations from typical patterns in brain structure. Their usefulness, however, is dependent on their applicability to diverse ethnoracial groups. With this in mind, we developed age- and sex-specific normative models for cortical thickness, surface area, and subcortical volumes using brain scans from 37,407 healthy individuals from a diverse international sample. Here we demonstrate the validity of these models in diverse and distinct populations. Specifically, we tested these pre-trained models on independent samples of healthy individuals that either self-identified as Black, South Asian, East Asian Chinese, East Asian Japanese, or we categorized as African, Admixed American, East Asian, and European based on their genetic ancestry. Regardless of ethnoracial definition, the performance of the pretrained models in these samples was exceptionally high; the relative mean absolute error for each regional brain morphometry measure was less than 10% across all the distinct ethnoracial groups. These findings affirm the broad applicability of our models, ensuring that brain morphometry assessments using these models are accurate and reliable for individuals regardless of background. This broad applicability has significant implications for advancing personalized medicine and improving health outcomes in diverse populations.

## INTRODUCTION

Normative modeling of neuroimaging-derived brain morphometry measures has the potential to inform about the clinical significance of deviations from typical neurostructural patterns^1-11^. As a result, several studies have recently emerged presenting different normative models for brain morphometry^12-18^.

The CentileBrain Project (www.centilebrain.org) is a major initiative of the Lifespan Working Group of the ENIGMA (Enhancing NeuroImaging Genetics through Meta Analysis) Consortium^19^ aiming to provide a robust and empirically validated platform of normative models of neuroimaging measures of brain organization. As part of this evolving initiative, we have developed race-neutral, sex-specific normative models of FreeSurfer-extracted regional measures of subcortical volumes, cortical thickness, and cortical surface area derived from an ethnoracial diverse sample of >37,000 healthy individuals from Australia, East Asia, Europe, North America and South Africa^13^. In the processes of developing the CentileBrain normative models of brain morphometry, our team undertook extensive benchmarking to ensure optimization of the algorithm and key covariates used and reliable model performance in individuals ranging from early childhood to late adulthood^13^. Importantly, our group also defined the minimal sample size requirements for robust and reproducible normative models^13^. The CentileBrain normative models are freely available through a dedicated platform (https://centilebrain.org/) and are being widely used by the neuroscience community with over 2,500 users from 64 countries.

In this study, we address the issue of ethnoracial diversity in the normative modeling of brain morphometry. Some neuroimaging studies have reported ethnoracial variation in brain morphometry^20-23^ but the literature is dominated by samples of non-white individuals^24^. Despite calls for increasing racial diversity in all medical research^25, 26^, there is no consensus on the definition of ethnoracial categories. While race and ethnicity correlate with genetic ancestry^27, 28^, ethnoracial labels based on physical attributes or historical context are broad, imprecise, and potentially harmful as they may perpetuate inequalities and misconceptions^29^. The appropriateness of existing ethnoracial adjustments in multiple diagnostic algorithms and practice guidelines used in medicine is currently being widely questioned, as such adjustments have been found to be unhelpful and potentially clinically detrimental^30^. For instance, in African American patients with chronic obstructive pulmonary disease, subjective symptoms and clinical signs align better when race-specific reference values are not used^31^.

As the CentileBrain morphometry models are race-neutral, as no adjustments were made for ethnoracial groupings, we tested their generalizability across distinct ethnoracial groups. To this end, we evaluated the performance of the race-neutral models on independent datasets where ethnoracial identity was designated as Black, South Asian, East Asian, Admixed American, or European White, based either on participants’ self-report or genetic ancestry.

## RESULTS

We applied the pre-trained, sex-specific and race neutral model to two sets of independent samples. The self-reported ethnoracial groups comprised individuals identifying as Black (N=284), South Asian (N=376), Japanese East Asian (N=970), and Chinese East Asian (N=1,136). Ancestry-defined ethnoracial groups comprised 207 individuals with African ancestry (AFR), 112 individuals with Admixed American ancestry (AMR), 54 individuals with East Asian ancestry (EAS), and 856 individuals with European (EUR) ancestry. The performance of each pre-trained model on the morphometric data of each distinct ethnoracial group was evaluated using the Mean Absolute Error (MAE) and the Root Mean Square Error (RMSE) which are the two most widely used measures of model accuracy. The comparative accuracy of the models as applied to the distinct ethnoracial samples was assessed by relative mean absolute error (RMAE) computed by dividing the MAE value by the true quantification of the regional morphometric measure.

The MAE and RMSE values obtained from the self-reported ethnoracial groups are shown in Supplementary Tables S1-S3, and Tables S7-S9. The MAE and RMSE values obtained from the genetic ancestry ethnoracial groups are shown in Supplementary Tables S4-S6, and Tables S10-S12. Regardless of whether ethnoracial status was defined by self-report or genetic ancestry, the correlation coefficients between the MAE or RMSE values of regional morphometric models in each ethnoracial group were highly correlated with those of the corresponding race-neutral CentileBrain models, with coefficients ranging from 0.85 to 0.99 (Figure 1 and Figure 2, Supplementary Figure S1-S6). Moreover, the mean MAE and RMSE across all regional morphometric models in each ethnoracial group did not exceed 0.4 standard deviation from the mean MAE and RMSE of the race-neutral CentileBrain models. For each ethnoracial group, the mean RMAE values for subcortical volume and surface area measures were below 10% across all brain regions, and for cortical thickness measures, the mean RMAE values across all brain regions were below 5% (Supplementary Tables S13-S18). We illustrate these findings using the left thalamic volume and left medial orbitofrontal cortical thickness and surface area as exemplars (Figure 3 and supplementary Figure S7). The normative deviation scores (Z-scores) in each ethnoracial group and in the race-neutral CentileBrain models were illustrated using the left thalamic volume and left medial orbitofrontal cortical thickness and surface area as exemplars (Figure 4 and supplementary Figure S8). The results showed that the average Z-score for each considered group was close to zero (Supplementary Table S19-S21), indicating the stability of applying the CentileBrain pre-trained models across diverse samples.

**Figure 1.**
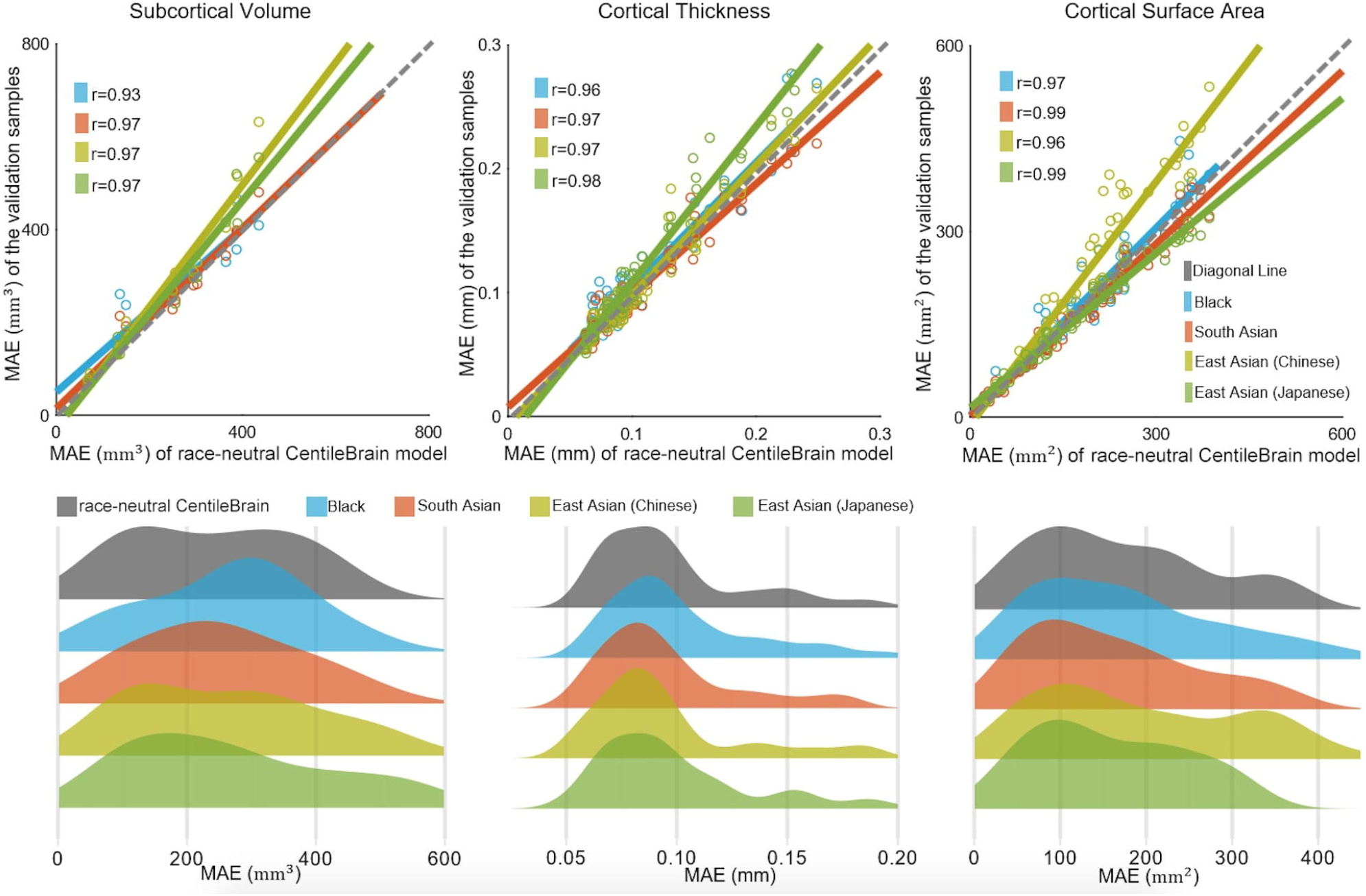
Top panel: scatterplot of the mean absolute error (MAE) values obtained from the self-reported ethnoracial groups and the corresponding values from the race-neutral CentileBrain model of female participants. Lower panel: distributions of the MAE values across the 14 subcortical volumes and cortical thickness and surface area across the 68 cortical regions for the race-neutral CentileBrain model and each ethnoracial group of female participants. Each circle denotes a regional morphometric measure. Different colors indicate different ethnoracial groups. The Pearson’s correlation coefficient (r) was computed between the MAE values of each ethnoracial group with those obtained in the race-neutral CentileBrain model. Additional information is provided in supplementary Table S1-S3. The results of RMSE were consistent with results of MAE, the results of RMSE are presented in supplementary Figure S1. In both sexes, the pattern identified was consistent for all region-specific models. The corresponding data for males are presented in supplementary Figure S2 and Figure S3.

**Figure 2.**
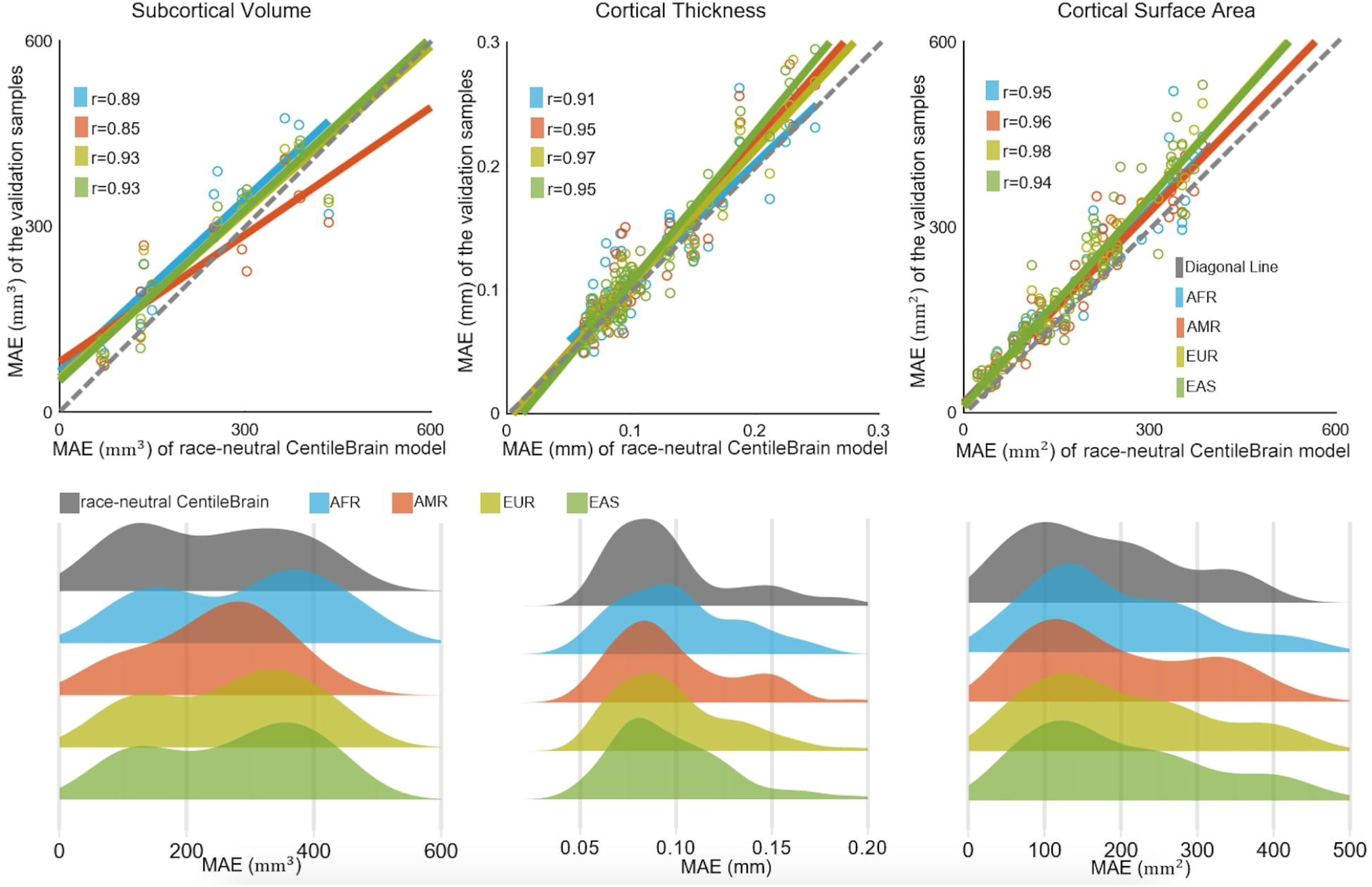
Top panel: scatterplot of the mean absolute error (MAE) values obtained from the genetic ancestry-determined ethnoracial groups and the corresponding values from the race-neutral CentileBrain model of female participants. Lower panel: distributions of the MAE values across the 14 subcortical volumes and cortical thickness and surface area across the 68 cortical regions for the race-neutral CentileBrain model and each ethnoracial group of female participants. Each circle denotes a regional morphometric measure. Different colors indicate different ethnoracial groups. The Pearson’s correlation coefficient (r) was computed between the MAE values of each ethnoracial group with those obtained in the race-neutral CentileBrain model. Additional information is provided in supplementary Table S4-S6. The results of RMSE were consistent with results of MAE, the results of RMSE are presented in supplementary Figure S4. In both sexes, the pattern identified was consistent for all region-specific models. The corresponding data for males are presented in supplementary Figure S5 and Figure S6. AFR=African; AMR=Admixed American; EAS=East Asian; EUR=European.

**Figure 3.**
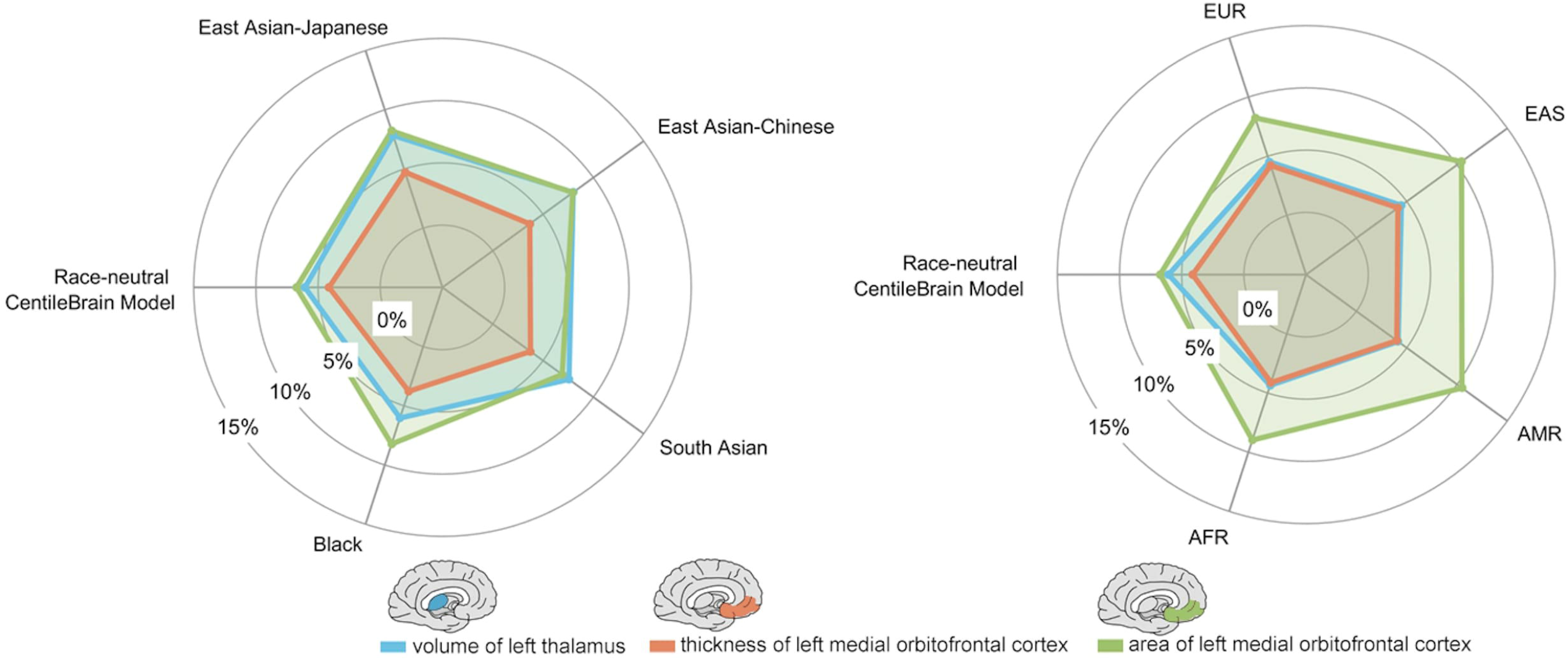
Left panel: spider plot of the relative mean absolute error (RMAE) values obtained from the self-reported ethnoracial groups and the corresponding values from the race-neutral CentileBrain model of female participants. Right panel: spider plot of the RMAE values obtained from the genetic ancestry-determined ethnoracial groups and the corresponding values from the race-neutral CentileBrain model of female participants. We illustrate these findings for females using the left thalamic volume and left medial orbitofrontal cortical thickness and surface area as exemplars (the corresponding data in males are in supplementary Figure S7). AFR=African; AMR=Admixed American; EAS=East Asian; EUR=European.

**Figure 4.**
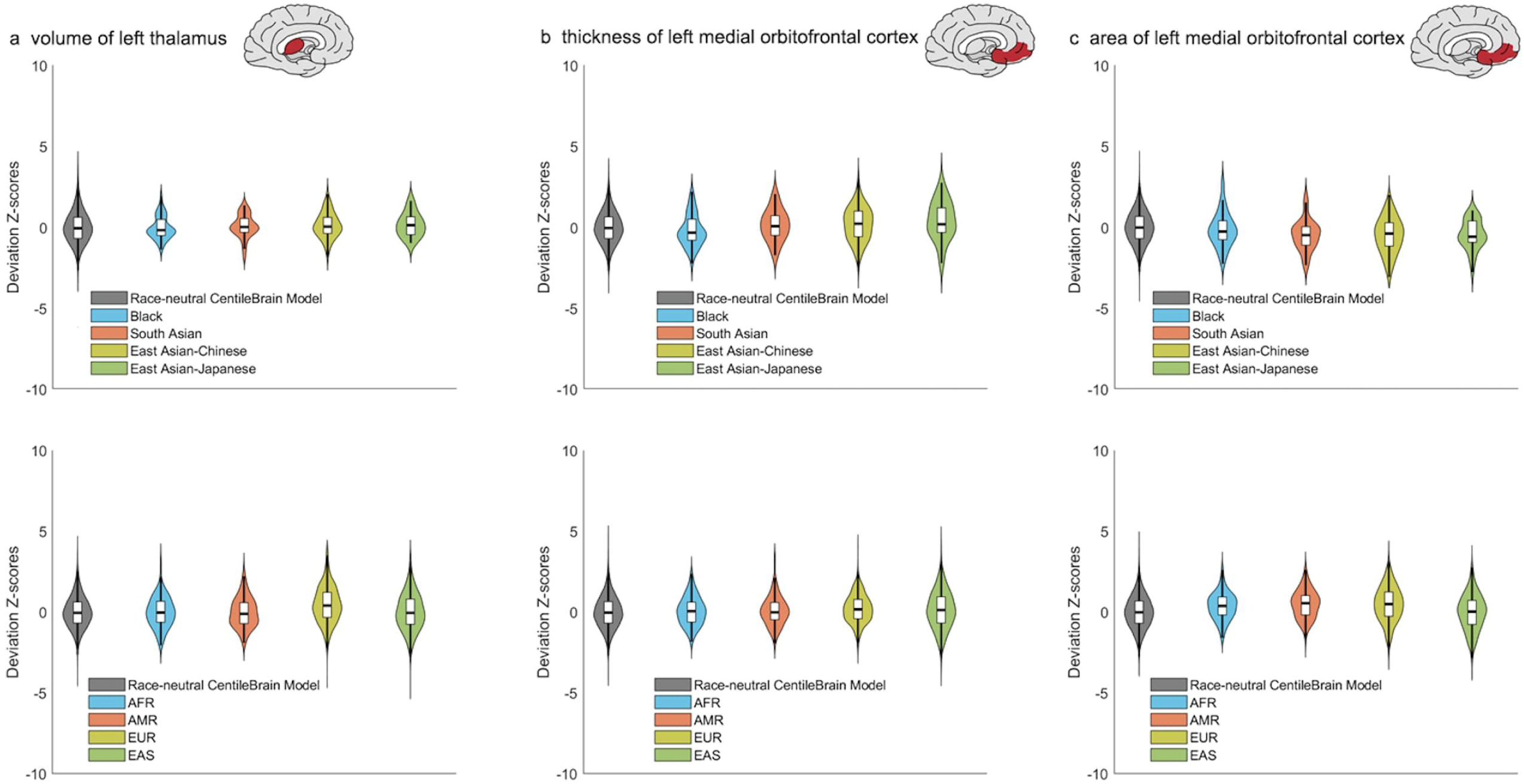
Normative deviation scores (Z-scores) in each ethnoracial group and in the race-neutral CentileBrain model of female participants. We illustrate these findings for females using the left thalamic volume (a) and left medial orbitofrontal cortical thickness (b) and surface area (c) as exemplars. The corresponding data in males are in supplementary Figure S8. AFR=African; AMR=Admixed American; EAS=East Asian; EUR=European.

## DISCUSSION

The present study provides empirical evidence that race-neutral normative models of brain morphometry, developed from a sample comprising a mixture of ethnoracial groups, can generalize to distinct ethnoracial samples.

It is widely acknowledged that neuroimaging research is significantly biased toward including mostly White participants^25^, although recent initiatives aim to redress this imbalance^32-35^. Normative modeling of brain morphometric data is a relatively recent addition to the statistical handling of neuroimaging data, with increasing uptake by research groups ^5,14, 36-40^, because it has the potential to inform about individual-level deviations from normative values given a person’s age and sex. This study addresses a key knowledge gap regarding the generalizability of race-neutral normative models to distinct ethnoracial groups. The results demonstrate that the accuracy of the race-neutral CentileBrain normative models of brain morphometry was similar across independent samples from diverse ethnoracial groups, regardless of how ethnoracial identity was defined. To our knowledge, the CentileBrain normative models are the only ones with empirical support for their generalizability to distinct ethnoracial groups.

To account for variations in brain volume, thickness, and area values across different ethnoracial groups compared to the model development sample, we introduced a relative MAE (RMAE) metric. This metric incorporates the variability of the true quantification of brain morphometric measures into the MAE measure. The results showed that, on average, the MAE value for each regional morphometric measure was less than 10% of the respective measure across all ethnoracial groups. Notably, the RMAE was particularly low especially for cortical thickness across all samples, with an average value smaller than 5%. These results underscore the high accuracy of the race-neutral CentileBrain models in predicting morphometric measures in distinct ethnoracial groups.

We acknowledge that the sample used for the development of the race-neutral CentileBrain models does not include all possible ethnoracial groups and that it included an over-representation of individuals self-identifying as White. This reflects the current state of the field; it would have been challenging to include equal representations of diverse ethnoracial groups while maintaining an adequate sample size for robust sex-specific modeling. We also acknowledge that the size of the independent ethnoracial samples is relatively modest, but we are strongly encouraged by the very robust RMAE values across all ethnoracial groups. We specifically avoided the inclusion of other explanatory variables, such as childhood adversity or socioeconomic status, which are known to influence brain morphometry^41,42^ as their inclusion in the models would reduce the ability to assess disparities arising from such exposures as was the case with the inclusion of race in other normative models in medicine.^30^ An alternate approach might have been to attempt to predict ethnoracial groups from brain morphometry reference data.^43^ We intentionally avoided this line of analysis because ethnoracial constructs are biologically imprecise, and using them as predicted outcomes could unintentionally imply that they represent biologically meaningful categories.

In conclusion, we present empirically validated evidence that the race-neutral CentileBrain normative models for brain morphometric measures can be effectively applied to samples from diverse ethnoracial backgrounds. Future research should continue to enhance the inclusivity and robustness of these models by incorporating a wider range of ethnoracial groups and additional relevant variables.

## METHODS

### Normative Models of Brain Morphometry

The CentileBrain Project recently released 150 normative models corresponding to FreeSurfer derived (http://surfer.nmr.mgh.harvard.edu/) regional measures of cortical thickness (N=68), cortical surface area (N=68) and regional subcortical volume (N=14) respectively based on the Desikan-Killiany Atlas^44^, and Aseg Atlas^45^. Details have been published^13^ and the relevant scripts and models are freely available through a dedicated web platform (www.centilebrain.org). Briefly, models were developed using multivariable fractional polynomial regression with the nonlinear fractional polynomials of age and linear global measures of the corresponding morphometric measure (i.e., intracranial volume, mean cortical thickness, and mean surface area). The model development sample consisted of 37,407 individuals (53.33% female) aged 3-90 years from 87 international datasets and comprised individuals self-identifying as White (95%), Black (3%), or Asian (2%). This sample was only used for the CentileBrain model development. The generalizability of the CentileBrain models to distinct ethnoracial groups was conducted in completely independent samples described below. As previously mentioned all the CentileBrain morphometry normative models are race-neutral and also sex-specific to accommodate sex differences in brain morphometry^46-48^.

The performance of each pre-trained model on the morphometric data of each distinct ethnoracial group was evaluated using the Mean Absolute Error (MAE) and the Root Mean Square Error (RMSE) which are the two most widely used measures of model accuracy. The MAE is the average absolute difference between the predicted and observed values of a variable of interest; it considers the magnitude of all errors (i.e., over- and under-estimates), it is robust to outliers and easy to interpret as it is expressed in the same units as the variable of interest. The RMSE is the square root of the average of squared differences between the predicted and observed values of a variable of interest and serves as a measure of the alignment between predicted and observed values.

### Samples

Data from six independent datasets were used to test the generalizability of the CentileBrain normative models of brain morphometry in individuals from distinct ethnoracial groups (Table 1). Depending on the available information, ethnoracial identity in some samples was based on participants’ self-report, while in others it was based on genetic ancestry. Although self-reported ethnoracial labels are biologically imprecise, they are still commonly used in most neuroimaging studies, as well as in medical research and practice, to encode ethnoracial information.

**Table 1.** Diverse Ethnoracial Datasets Used for Model Generalizability.

| Ethnoracial Group | Dataset Abbreviation | Scanner Vendor | Scanner Strength (Tesla) | FreeSurfer Version | Age (years) Mean | Age (years) Range | Male/Female N |
| --- | --- | --- | --- | --- | --- | --- | --- |
| **Self-reported Groups** |  |  |  |  |  |  |  |
| Black | UK BIOBANK<sup>1</sup> | Siemens Skyra | 3T | 7.1 | 59.22(7.28) | 47-79 | 125/159 |
| South Asian | UK BIOBANK | Siemens Skyra | 3T | 7.1 | 60.34(8.34) | 44-79 | 237/139 |
| East Asian (Chinese) | UK BIOBANK | Siemens Skyra | 3T | 7.1 | 59.86(6.97) | 47-76 | 46/77 |
|  | SALD<sup>2</sup> | Siemens TIM Trio | 3T | 7.1 | 45.16(17.45) | 19-80 | 185/307 |
|  | SLIM<sup>3</sup> | Siemens TIM Trio | 3T | 5.3 | 20.06(1.28) | 17-27 | 229/291 |
|  | CHCP<sup>4</sup> | Siemens Prisma | 3T | 7.1 | 34.14(18.19) | 18-79 | 166/195 |
| East Asian (Japanese) | COCORO<sup>5</sup> | General Electric Signa EXCITE; General Electric Discovery MR750 | 1.5T; 3T | 5.3 | 34.13(14.25) | 18-75 | 481/489 |
| Total | - | - | - | - | 39.97(18.66) | 17-80 | 1,468/1,657 |
| **Genetic Ancestry Groups** |  |  |  |  |  |  |  |
| African (AFR) | ABCD<sup>6</sup> | Siemens Prisma VE11B-C; Philips Achieva dStream; Philips Ingenia; GE Discovery MR750; GE DV25-26 | 3 | 5.3 | 9.91(0.60) | 8.91-10.91 | 50/54 |
| Admixed American (AMR) | ABCD | Siemens Prisma VE11B-C; Philips Achieva dStream; Philips Ingenia; GE Discovery MR750; GE DV25-26 | 3 | 5.3 | 9.91(0.56) | 8.91-10.99 | 26/31 |
| East Asian (EAS) | ABCD | Siemens Prisma VE11B-C; Philips Achieva dStream; Philips Ingenia; GE Discovery MR750; GE DV25-26 | 3 | 5.3 | 9.91(0.66) | 8.91-10.91 | 21/33 |
| European (EUR) | ABCD | Siemens Prisma VE11B-C; Philips Achieva dStream; Philips Ingenia; GE Discovery MR750; GE DV25-26 | 3 | 5.3 | 9.95(0.63) | 8.99-10.99 | 224/204 |
| Total | - | - | - | - | 9.94(0.62) | 8.91-10.99 | 321/322 |

ABCD=Adolescent Brain Cognitive Development; CHCP=Chinese Human Connectome Project; COCORO=Cognitive Genetics Collaborative Research Organization; SALD=Southwest University Adult Lifespan Dataset; SLIM=Southwest University Longitudinal Imaging Multimodal Study
1. https://www.ukbiobank.ac.uk/
2. Wei, D., et al. (2018). Structural and functional brain scans from the cross-sectional Southwest University adult lifespan dataset. Scientific Data, 5(1), 1-10.
3. Liu, W., et al. (2017). Longitudinal test-retest neuroimaging data from healthy young adults in southwest China. Scientific Data, 4(1), 1-9.
4. Ge, J., et al. (2023). Increasing diversity in connectomics with the Chinese Human Connectome Project. Nature Neuroscience, 26, 163-172.
5. Koshiyama, D., et al. (2022). Neuroimaging studies within Cognitive Genetics Collaborative Research Organization aiming to replicate and extend works of ENIGMA. Human Brain Mapping, 43(1), 182-193.
6. https://abcdstudy.org/

#### Samples with Self-reported Ethnoracial Identities

We used data from participants identifying as Black (N=284), South Asian (N=376), Japanese East Asian (N=970), and Chinese East Asian (N=1,136). The Chinese East Asian sample pooled participants from the Southwest University Adult Lifespan Dataset (SALD)^49^ and the Southwest University Longitudinal Imaging Multimodal Study (SLIM)^50^ and the UK Biobank (UKB) Study (https://www.ukbiobank.ac.uk)^51^. The Japanese East Asian sample included data from participants of the study of the Cognitive Genetics Collaborative Research Organization (COCORO).^32^ The samples comprising exclusively of participants that self-identified as Black or South Asian were derived from the UK Biobank Study.

#### Genetic ancestry Samples

Ethnoracial groups defined by genetic ancestry were derived from the Adolescent Brain Cognitive Development Study (ABCD; https://abcdstudy.org/)^52^ where each participant’s ancestry was determined using the “fastSTRUCTURE” algorithm^53^ with four ancestry factors corresponding to 4 super-populations: African ancestry (AFR), Admixed American Ancestry (AMR), East Asian (EAS), and European (EUR) ancestry. Among 11,875 participants from the ABCD dataset, we used data from 3,759 participants based on the absence of any lifetime psychiatric, neurological or medical morbidity. We further selected participants if they could be exclusively allocated to only one of the four super-population groups based on their estimated posterior probability of the factor for that group being higher than 0.99. Of the 3,759 ABCD participants, 1,230 met this criterion and comprised 207 individuals designated as AFR, 113 as AMR, 54 as EAS and 856 as EUR. We randomly selected half of these participants as generalizability validation samples (Table 1), and the remaining were used in the CentileBrain model development sample (Ge et al., 2024).

Aligned with the processes used for the CentileBrain normative model development, the standard pipelines in the FreeSurfer image analysis suite (http://surfer.nmr.mgh.harvard.edu/) were applied to the whole-brain T1-weighted images of the participants in the distinct ethnoracial groups to yield regional measures of cortical thickness and cortical surface area and regional subcortical volumes.

### Statistical Analysis

The CentileBrain race-neutral and sex-specific models for regional cortical thickness, cortical surface area, and subcortical volume measures were applied to the corresponding neuroimaging measures of each participant in each of the distinct ethnoracial groups. The accuracy of these models was assessed using the MAE and the RSME. Pearson correlation coefficient was used to examine the association between these values and the MAE and RSME of the corresponding race-neutral Centilebrain model. Further, the accuracy of the models as applied to the distinct ethnoracial samples was assessed by relative mean absolute error (RMAE) computed by dividing the MAE value by the true quantification of the regional morphometric measure. RMAE represents the ratio of the error between the observed and the predicted value of a variable of interest set to the measured value of all the observations. The advantage of this additional measure is that it provides a normalized estimate of prediction accuracy which helps in identifying models that perform well relative to the magnitude of the values being predicted, offering a clearer understanding of their predictive capability.

## DATA AVAILABILITY

The pre-trained CentileBrain Models are freely available at https://centilebrain.org/. The ABCD dataset can be accessed through the US National Data Archive (https://nda.nih.gov/); the CHCP dataset can be accessed through the Science Data Bank website (https://doi.org/10.11922/sciencedb.01374); the COCORO dataset is available upon reasonable request; the SALD and SLIM datasets can be accessed through the International Data-sharing Initiative (https://fcon_1000.projects.nitrc.org/indi/retro/sald.html and https://fcon_1000.projects.nitrc.org/indi/retro/southwestuni_qiu_index.html); and the UKB dataset can be access through the UK Biobank data-access protocol (https://www.ukbiobank.ac.uk/enable-your-research/apply-for-access).

## ACKNOWLEDGEMENT

Analyses were conducted using high performance computing (https://arc.ubc.ca/ubc-arc-sockeye) available through the Advanced Research Computing service, University of British Columbia. Some computations were performed at the Research Center for Computational Science, Okazaki, Japan (projects: NIPS, 15-IMS-C137, 16-IMS-C135, 17-IMS-C152, 18-IMS-C162, 19-IMS-C181, 20-IMS-C162, 21-IMS-C179, 22-IMS-C195).This research was supported by AMED under Grant Number JP21uk1024002 and JP24dk0307132, the Intramural Research Grant (6-1) for Neurological and Psychiatric Disorders of NCNP, and JSPS Grant-in-Aid for Scientific Research (C) JP23K07001.

## AUTHOR CONTRIBUTIONS

All authors contributed data and participated in the analyses and manuscript preparation. RG had overall oversight on data collation and data analyses and SF oversaw all aspects of the study and provided the first draft of the manuscript.

