## Supplement for "Generalizability of Normative Models of Brain Morphometry Across Distinct Ethnoracial Groups": CentileBrain Validation Supplement.pdf

**Supplementary Material**

### Supplementary Results

**Supplementary Figure S1**. Scatterplot and distribution plot of the RMSE values obtained from the self-reported ethnoracial groups and the corresponding values from the race-neutral CentileBrain model of female participants.
**Supplementary Figure S2**. Scatterplot and distribution plot of the MAE values obtained from the self-reported ethnoracial groups and the corresponding values from the race-neutral CentileBrain model of male participants.
**Supplementary Figure S3**. Scatterplot and distribution plot of the RMSE values obtained from the self-reported ethnoracial groups and the corresponding values from the race-neutral CentileBrain model of male participants.
**Supplementary Figure S4**. Scatterplot and distribution plot of the RMSE values obtained from the genetic ancestry-determined ethnoracial groups and the corresponding values from the race-neutral CentileBrain model of female participants.
**Supplementary Figure S5**. Scatterplot and distribution plot of the MAE values obtained from the genetic ancestry-determined ethnoracial groups and the corresponding values from the race-neutral CentileBrain model of male participants.
**Supplementary Figure S6**. Scatterplot and distribution plot of the RMSE values obtained from the genetic ancestry-determined ethnoracial groups and the corresponding values from the race-neutral CentileBrain model of male participants.
**Supplementary Figure S7.** Spider plot of the relative mean absolute error (RMAE) values obtained from the self-reported ethnoracial groups, genetic ancestry-determined ethnoracial groups, and corresponding values from the race-neutral CentileBrain model of male participants.
**Supplementary Figure S8.** Normative deviation scores (Z-scores) in each ethnoracial group and in the race-neutral CentileBrain model of male participants.

**Table S1**. Mean absolute error (MAE) and root-mean-square error (RMSE) values of each regional subcortical volume in self-reported ethnoracial groups and in the race-neutral CentileBrain model of female participants

**Table S2**. Mean absolute error (MAE) and root-mean-square error (RMSE) values of each cortical thickness in self-reported ethnoracial groups and in the race-neutral CentileBrain model of female participants

**Table S3**. Mean absolute error (MAE) and root-mean-square error (RMSE) values of each cortical surface area in self-reported ethnoracial groups and in the race-neutral CentileBrain model of female participants

**Table S4**. Mean absolute error (MAE) and root-mean-square error (RMSE) values of each regional subcortical volume in genetically determined ethnoracial groups and in the race-neutral CentileBrain model of female participants

**Table S5**. Mean absolute error (MAE) and root-mean-square error (RMSE) values of each cortical thickness in genetically determined ethnoracial groups and in the race-neutral CentileBrain model of female participants

**Table S6**. Mean absolute error (MAE) and root-mean-square error (RMSE) values of each cortical surface area in genetically determined ethnoracial groups and in the race-neutral CentileBrain model of female participants

**Table S7**. Mean absolute error (MAE) and root-mean-square error (RMSE) values of each regional subcortical volume in self-reported ethnoracial groups and in the race-neutral CentileBrain model of male participants

**Table S8**. Mean absolute error (MAE) and root-mean-square error (RMSE) values of each cortical thickness in self-reported ethnoracial groups and in the race-neutral CentileBrain model of male participants

**Table S9**. Mean absolute error (MAE) and root-mean-square error (RMSE) values of each cortical surface area in self-reported ethnoracial groups and in the race-neutral CentileBrain model of male participants

**Table S10**. Mean absolute error (MAE) and root-mean-square error (RMSE) values of each regional subcortical volume in genetically determined ethnoracial groups and in the race-neutral CentileBrain model of male participants

**Table S11**. Mean absolute error (MAE) and root-mean-square error (RMSE) values of each cortical thickness in genetically determined ethnoracial groups and in the race-neutral CentileBrain model of male participants

**Table S12**. Mean absolute error (MAE) and (RMSE) values of each cortical surface area in genetically determined ethnoracial groups and in the race-neutral CentileBrain model of male participants

**Table S13**. Relative mean absolute error (RMAE) values of each regional subcortical volume in ethnoracial groups and in the race-neutral CentileBrain model of female participants

**Table S14**. Relative mean absolute error (RMAE) values of each regional cortical thickness in ethnoracial groups and in the race-neutral CentileBrain model of female participants

**Table S15**. Relative mean absolute error (RMAE) values of each regional surface area in ethnoracial groups and in the race-neutral CentileBrain model of female participants

**Table S16**. Relative mean absolute error (RMAE) values of each regional subcortical volume in ethnoracial groups and in the race-neutral CentileBrain model of male participants

**Table S17**. Relative mean absolute error (RMAE) values of each regional cortical thickness in ethnoracial groups and in the race-neutral CentileBrain model of male participants

**Table S18**. Relative mean absolute error (RMAE) values of each regional surface area in ethnoracial groups and in the race-neutral CentileBrain model of male participants

**Table S19**. Average deviation Z-scores (SD) of subcortical volume measure in each ethnoracial group and in the race-neutral CentileBrain models

**Table S20**. Average deviation Z-scores (SD) of cortical thickness measure in each ethnoracial group and in the race-neutral CentileBrain models

**Table S21**. Average deviation Z-scores (SD) of surface area measure in each ethnoracial group and in the race-neutral CentileBrain models

### Supplementary Figures

**Figure S1.** Top panel: scatterplot of the root-mean-square error (RMSE) values obtained from the self-reported ethnoracial groups and the corresponding values from the model development sample of female participants. Lower panel: distributions of the RMSE values across the 14 subcortical volumes and cortical thickness and surface area across the 68 cortical regions for the model development sample and each ethnoracial group of female participants.

Each circle denotes a regional morphometric measure. Different colors indicate different ethnoracial groups. The Pearson's correlation coefficient (r) was computed between the RMSE values of each ethnoracial group with those obtained in the model development sample.

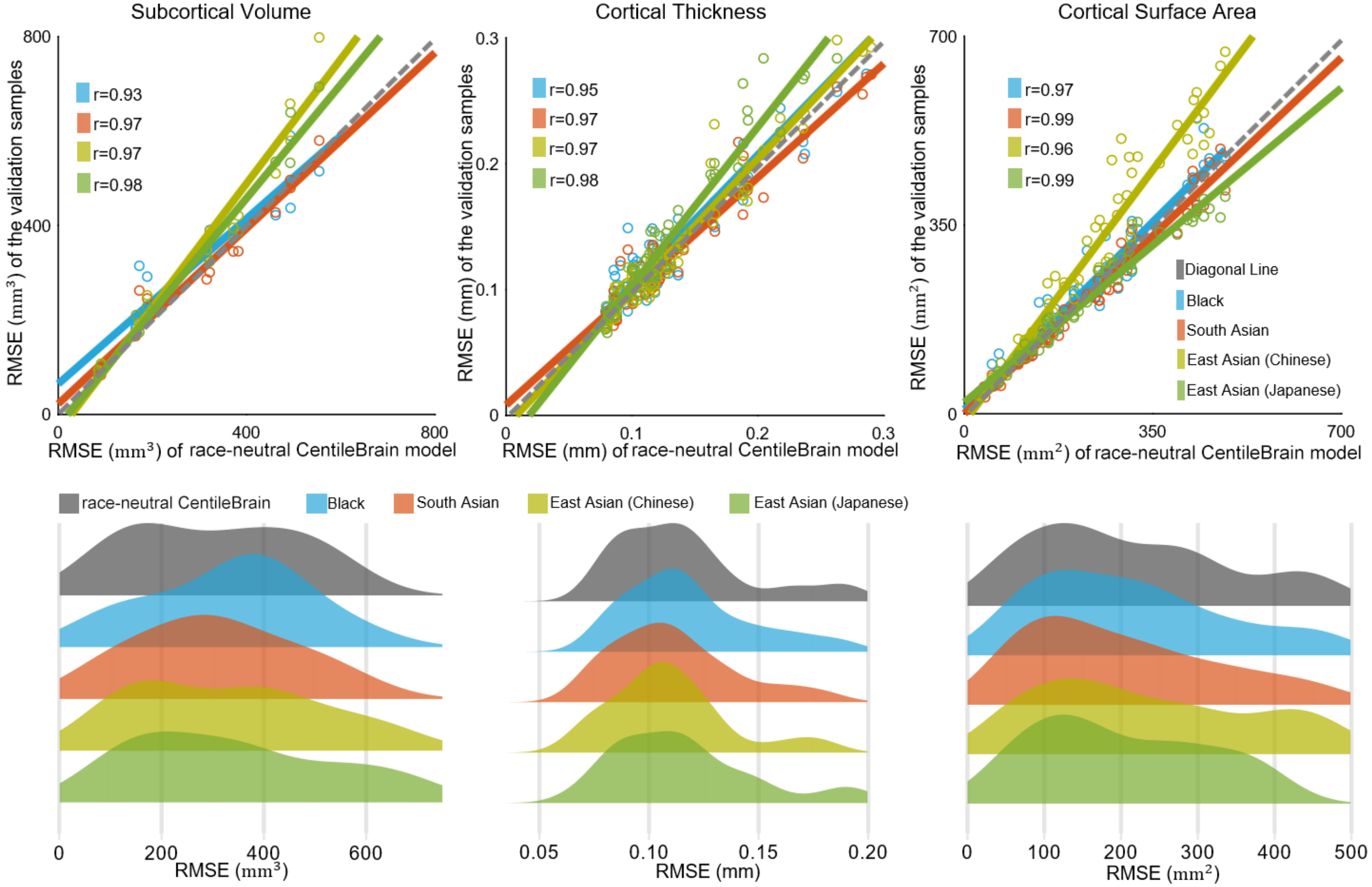

**Figure S2.** Top panel: scatterplot of the mean absolute error (MAE) values obtained from the self-reported ethnoracial groups and the corresponding values from the model development sample of male participants. Lower panel: distributions of the MAE values across the 14 subcortical volumes and cortical thickness and surface area across the 68 cortical regions for the model development sample and each ethnoracial group of male participants.

Each circle denotes a regional morphometric measure. Different colors indicate different ethnoracial groups. The Pearson's correlation coefficient (r) was computed between the MAE values of each ethnoracial group with those obtained in the model development sample.

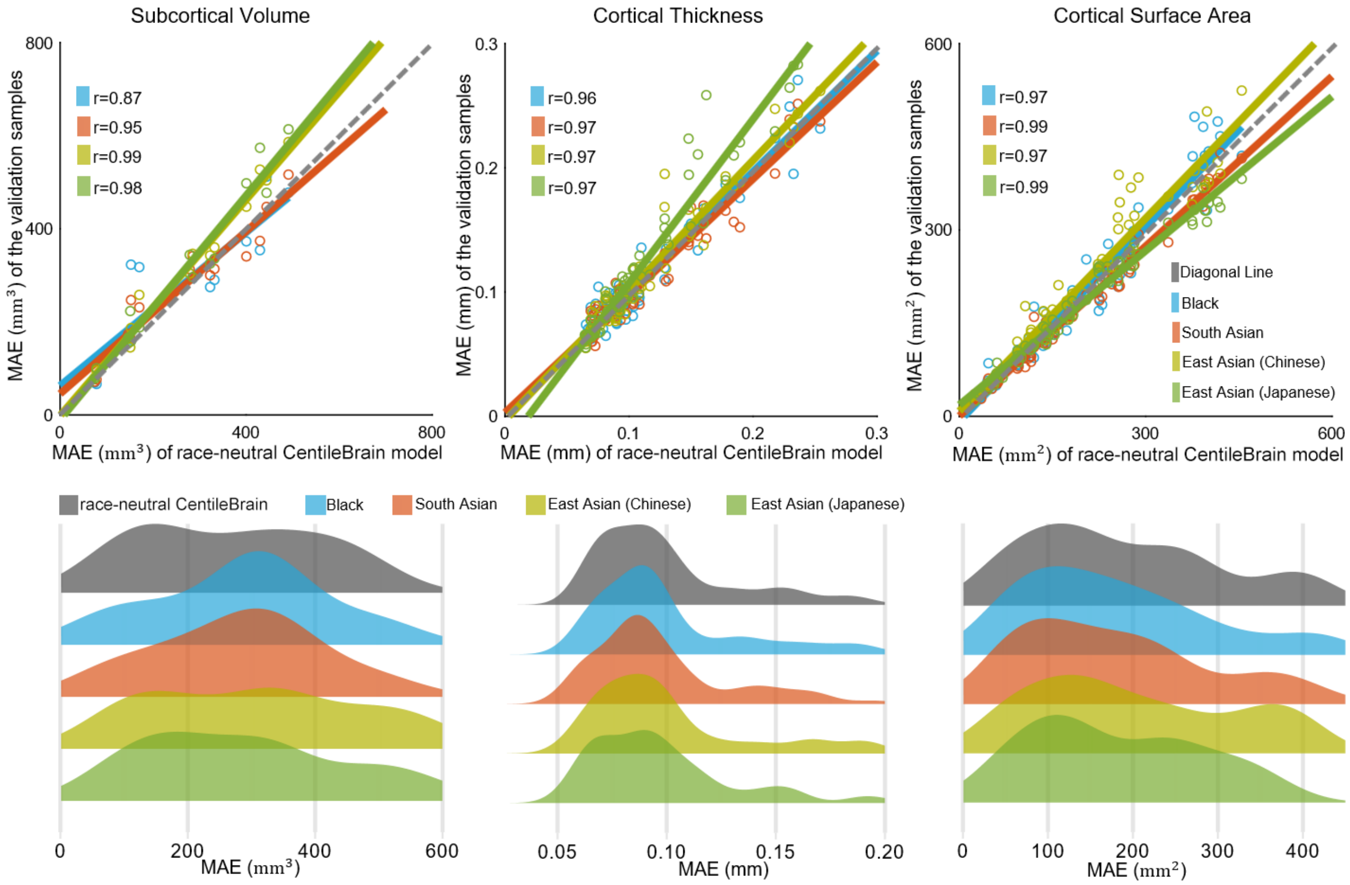

**Figure S3.** Top panel: scatterplot of the root-mean-square error (RMSE) values obtained from the self-reported ethnoracial groups and the corresponding values from the model development sample of male participants. Lower panel: distributions of the RMSE values across the 14 subcortical volumes and cortical thickness and surface area across the 68 cortical regions for the model development sample and each ethnoracial group of male participants.

Each circle denotes a regional morphometric measure. Different colors indicate different ethnoracial groups. The Pearson's correlation coefficient (r) was computed between the RMSE values of each ethnoracial group with those obtained in the model development sample.

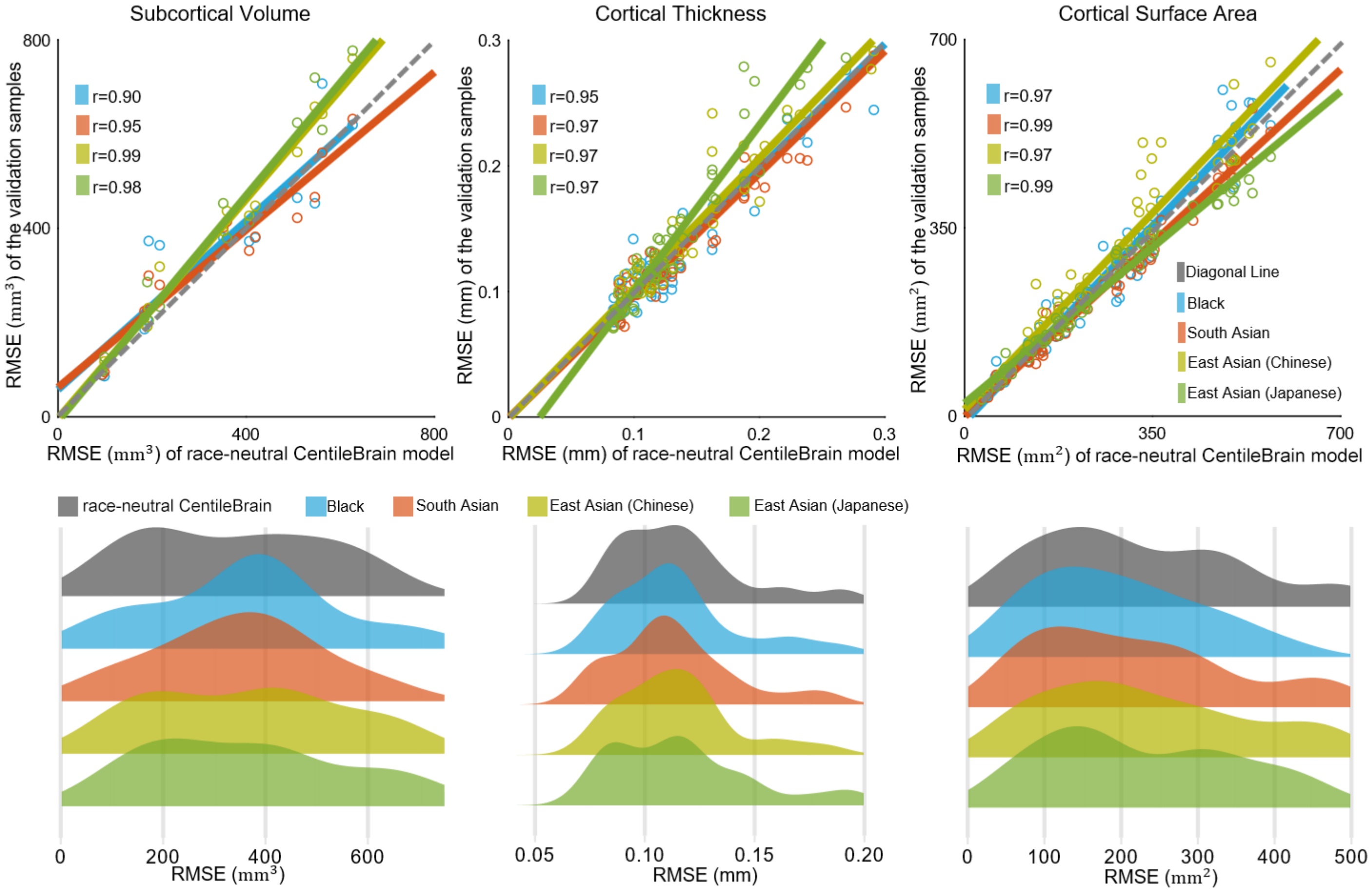

**Figure S4.** Top panel: scatterplot of the root-mean-square error (RMSE) values obtained from the genetic ancestry-determined ethnoracial groups and the corresponding values from the model development sample of female participants. Lower panel: distributions of the RMSE values across the 14 subcortical volumes and cortical thickness and surface area across the 68 cortical regions for the model development sample and each ethnoracial group of female participants.

Each circle denotes a regional morphometric measure. Different colors indicate different ethnoracial groups. The Pearson's correlation coefficient (r) was computed between the RMSE values of each ethnoracial group with those obtained in the model development sample.

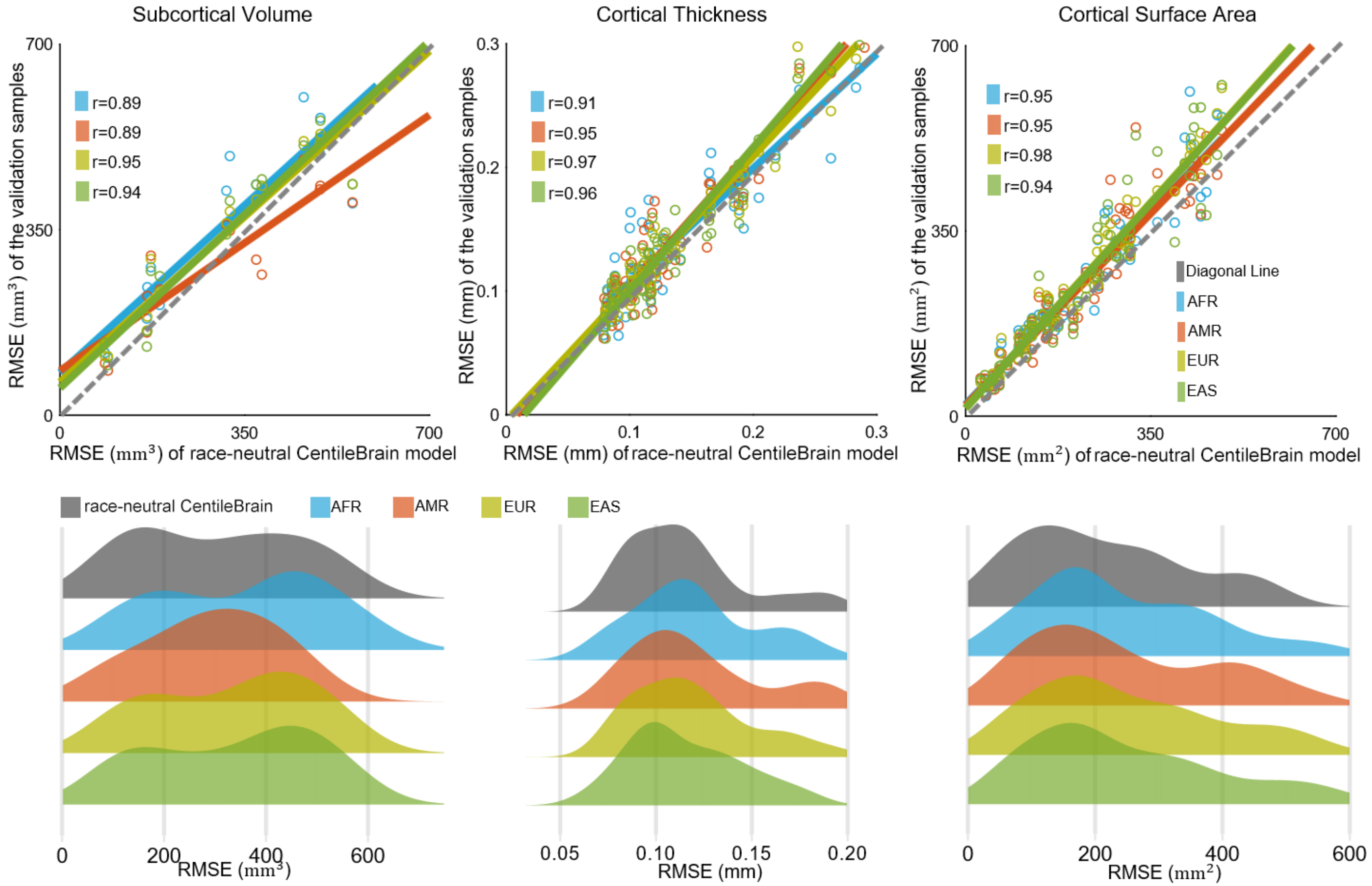

**Figure S5.** Top panel: scatterplot of the mean absolute error (MAE) values obtained from the genetic ancestry-determined ethnoracial groups and the corresponding values from the model development sample of male participants. Lower panel: distributions of the MAE values across the 14 subcortical volumes and cortical thickness and surface area across the 68 cortical regions for the model development sample and each ethnoracial group of male participants.

Each circle denotes a regional morphometric measure. Different colors indicate different ethnoracial groups. The Pearson's correlation coefficient (r) was computed between the MAE values of each ethnoracial group with those obtained in the model development sample.

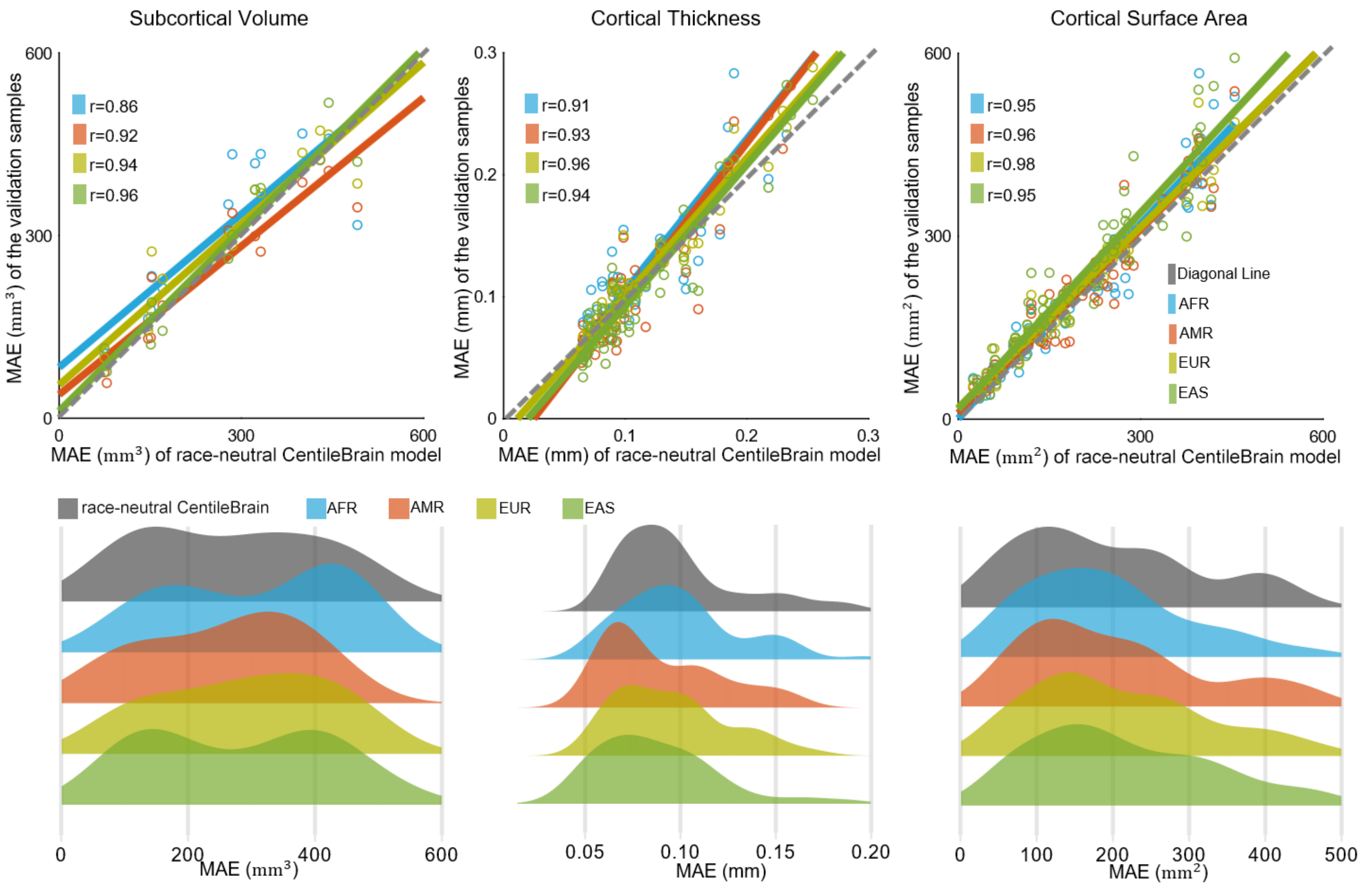

**Figure S6.** Top panel: scatterplot of the root-mean-square error (RMSE) values obtained from the genetic ancestry-determined ethnoracial groups and the corresponding values from the model development sample of male participants. Lower panel: distributions of the RMSE values across the 14 subcortical volumes and cortical thickness and surface area across the 68 cortical regions for the model development sample and each ethnoracial group of male participants.

Each circle denotes a regional morphometric measure. Different colors indicate different ethnoracial groups. The Pearson's correlation coefficient (r) was computed between the RMSE values of each ethnoracial group with those obtained in the model development sample.

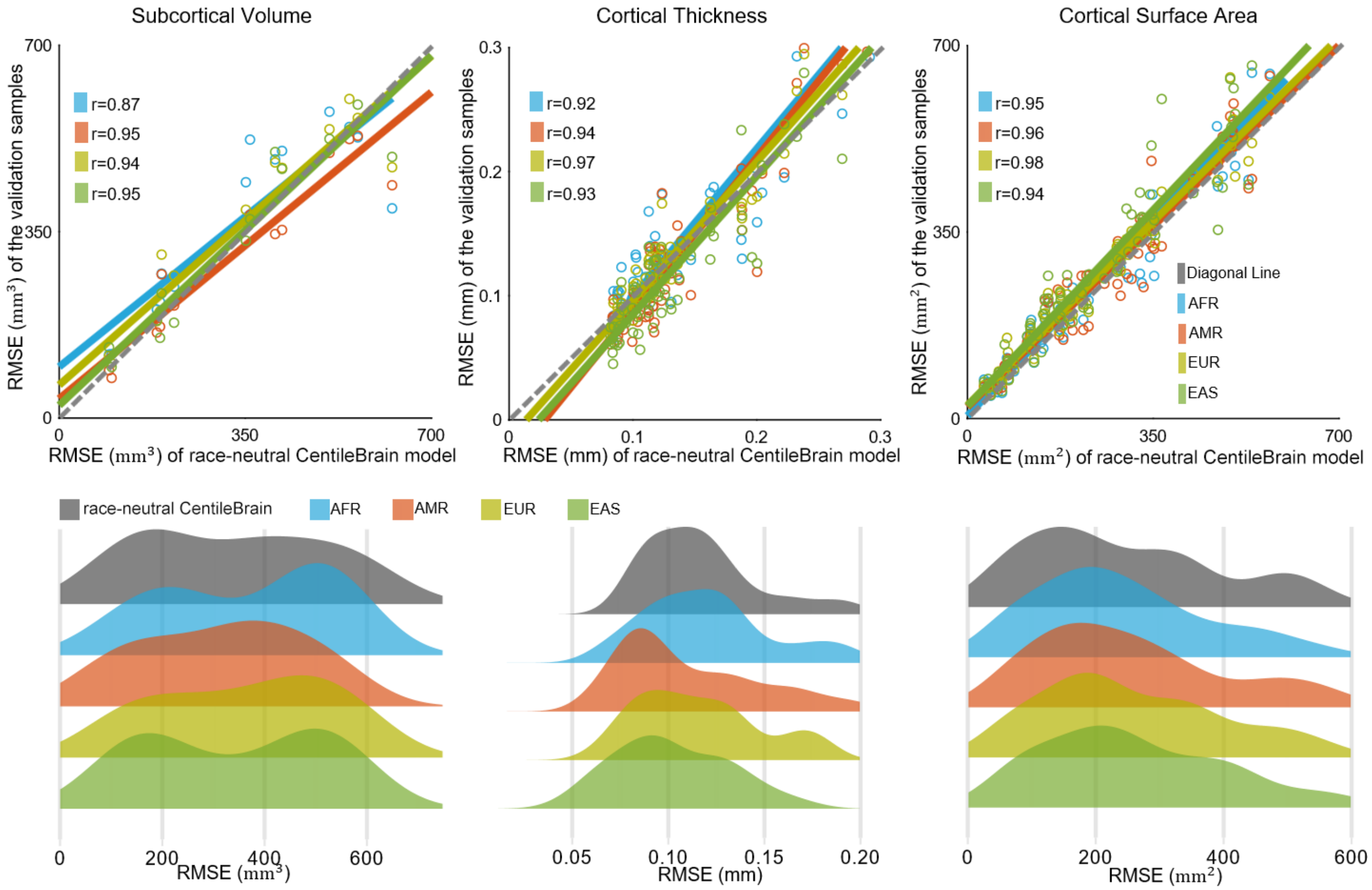

**Figure S7.** Left panel: spider plot of the relative mean absolute error (RMAE) values obtained from the self-reported ethnoracial groups and the corresponding values from the race-neutral CentileBrain model of male participants. Right panel: spider plot of the RMAE values obtained from the genetic ancestry-determined ethnoracial groups and the corresponding values from the race-neutral CentileBrain model of male participants. We illustrate these findings for males using the left thalamic volume and left medial orbitofrontal cortical thickness and surface area as exemplars. AFR=African; AMR=Admixed American; EAS=East Asian; EUR=European.

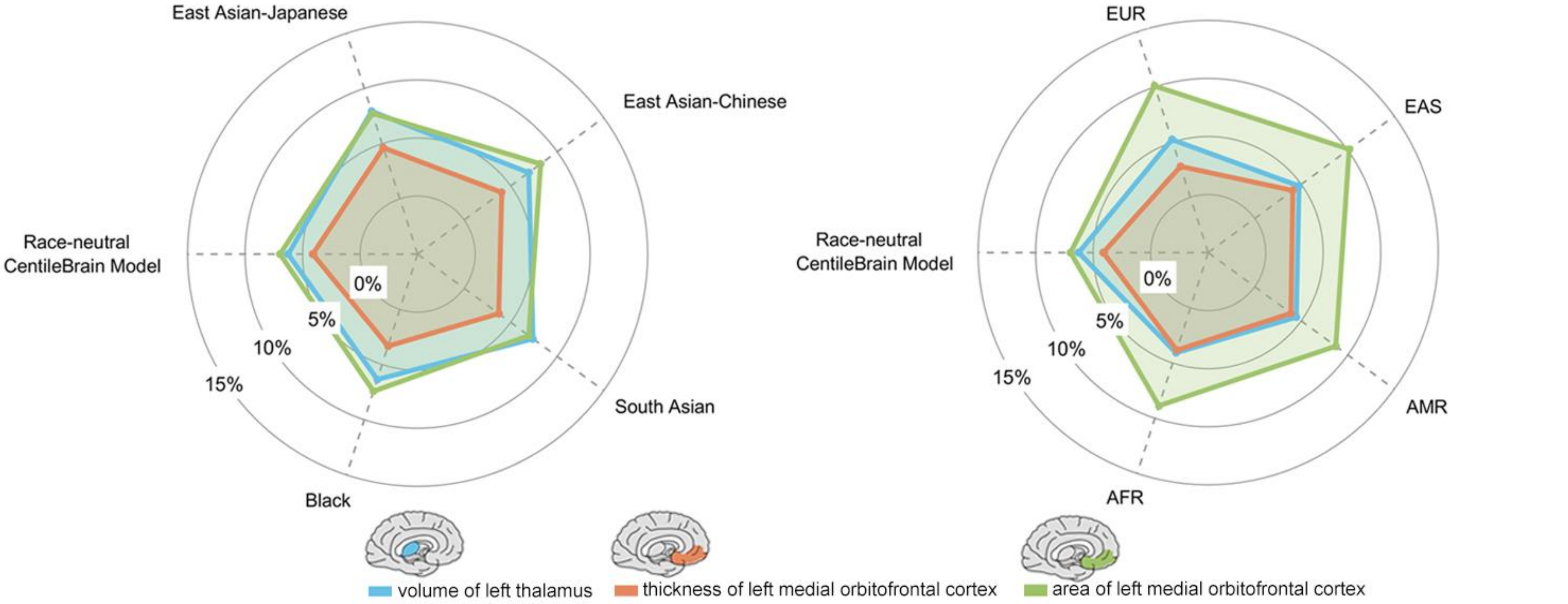

**Figure S8.** Normative deviation scores (Z-scores) in each ethnoracial group and in the race-neutral CentileBrain model of male participants. We illustrate these findings for males using the left thalamic volume (a) and left medial orbitofrontal cortical thickness (b) and surface area (c) as exemplars.
AFR=African; AMR=Admixed American; EAS=East Asian; EUR=European.

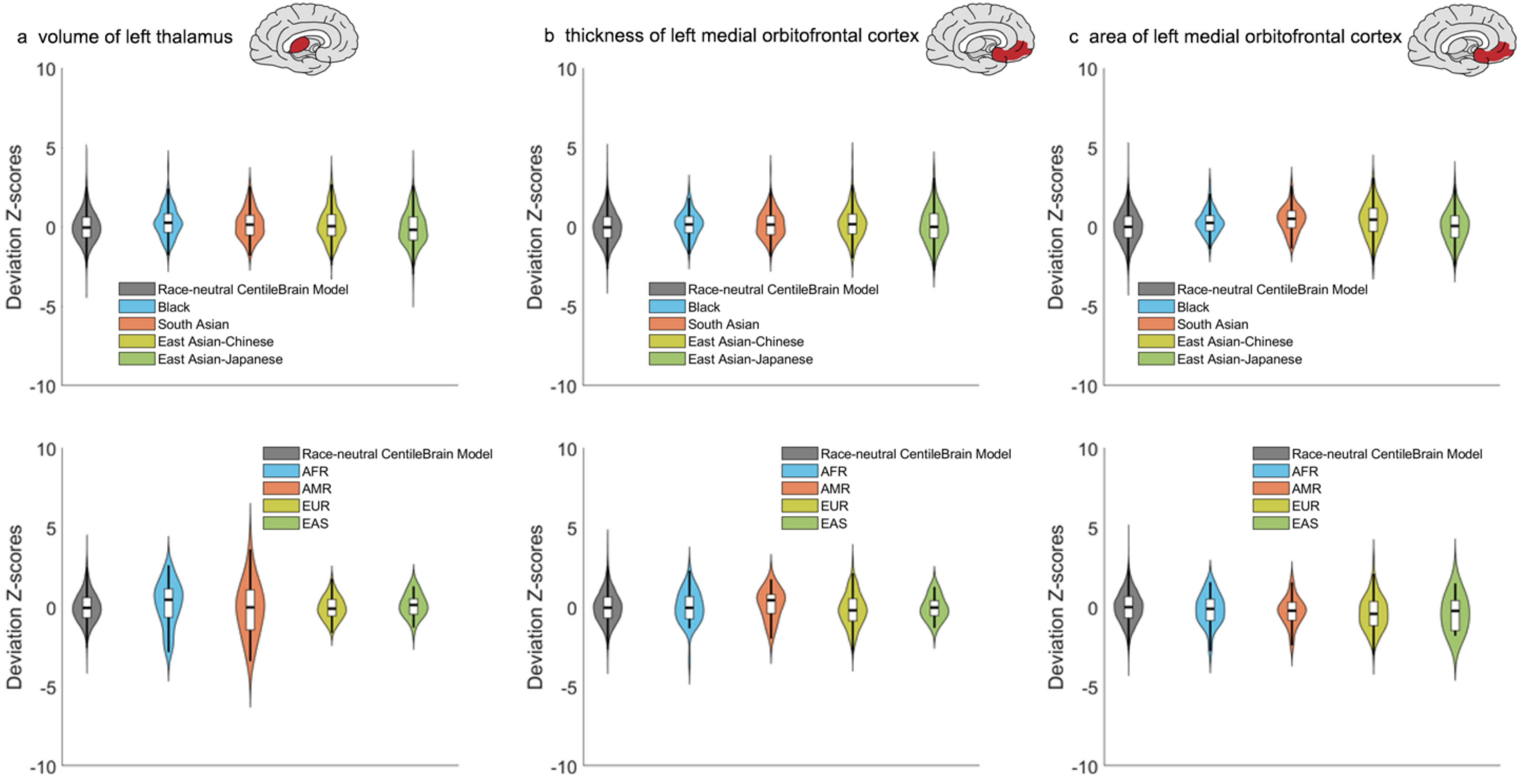
